# Divergent proinflammatory immune responses associated with the differential susceptibility of cattle breeds to Tuberculosis

**DOI:** 10.1101/2023.03.30.534906

**Authors:** Rishi Kumar, Sripratyusha Gandham, Avi Rana, Hemant Kumar Maity, Uttam Sarkar, Bappaditya Dey

**Affiliations:** National Institute of Animal Biotechnology, Hyderabad, Telangana, India, PIN 500032; Regional Centre for Bioinformatics, Faridabad, Haryana, India, PIN 121001; West Bengal University of Animal and Fishery Sciences. Kolkata, West Bengal, India, PIN 700037

**Author notes:** **Corresponding Author:** Dr. Bappaditya Dey Scientist-E, National Institute of Animal Biotechnology, Hyderabad, Telangana, India - 500032, Email ID.

**Keywords:** Tuberculosis, Bovine tuberculosis, *Mycobacterium tuberculosis*, *Mycobacterium bovis*, BCG

## Abstract

Tuberculosis (TB) in the bovine is one of the most predominant chronic debilitating infectious diseases primarily caused by *Mycobacterium bovis*. Besides, the incidence of TB in humans due to *M. bovis*, and that in bovines due to *M. tuberculosis*-indicates cattle as a major reservoir of zoonotic TB. While India accounts for the highest global burden of both TB and multidrug-resistant TB in humans, systematic evaluation of bovine TB (bTB) prevalence in India is largely lacking. Recent reports emphasized markedly greater bTB prevalence in exotic and crossbred cattle compared to indigenous cattle breeds that represent more than one-third of the total cattle population in India, which is the largest globally. This study aimed at elucidating the immune responses underlying the differential bTB incidence in prominent indigenous (Sahiwal), and crossbred (Sahiwal x Holstein Friesian) cattle reared in India. Employing the standard Single Intradermal Tuberculin Test (SITT), and mycobacterial gene-targeting single as well as multiplex-PCR-based screening revealed higher incidences of bovine tuberculin reactors as well as *Mycobacterium tuberculosis* Complex specific PCR positivity amongst the crossbred cattle. Further, *ex vivo* mycobacterial infection in cultures of bovine peripheral blood mononuclear cells (PBMC) from SITT, and myco-PCR negative healthy cattle exhibited significantly higher intracellular growth of *M. bovis* BCG, and *M. tuberculosis* H37Ra in the crossbred cattle PBMCs compared to native cattle. In addition, native cattle PBMCs induced higher pro-inflammatory cytokines and signaling pathways, such as interferon-gamma (IFN-*γ*), interleukin-17 (IL-17), and tank binding kinase-1 (TBK-1) upon exposure to live mycobacterial infection in comparison to PBMCs from crossbred cattle that exhibited higher expression of IL-1β transcripts. Together, these findings highlight that differences in the innate immune responses of these cattle breeds might be contributing to the differential susceptibility to TB infection, and the resultant disparity in bTB incidence amongst indigenous, and crossbred cattle.

## Introduction

Bovine tuberculosis (bTB) is a globally prevalent chronic debilitating infectious disease of cattle with a considerable impact on the public health and farm economy. Of the 188 countries and territories globally reporting their bovine TB (bTB) situation to the World Organisation of Animal Health (OIE), 82 countries (44%) reported bTB prevalence [1]. Notably, while 97.6% of the affected countries reported bTB prevalence in livestock, 35.4% of countries documented bTB presence in both livestock and wildlife animals [1]. In addition, the incidence of TB in humans and bovines due to either the human or bovine tubercle bacilli signifies the impact of TB on livestock farming, and highlights its transmission between cattle and humans [2-4]. Since, about 54.6% of the total workforce in India is engaged in agriculture and animal husbandry, and livestock provides livelihood to two third of the rural community, therefore, bTB exerts a hugely adverse effect on public health [3, 5].

Bovine TB has been largely controlled in many high-income countries due to strict implementation of bTB control programs and policies, whereas in lower, and lower-middle-income countries, control of bTB still poses a major challenge [6, 7]. This is largely because of unhygienic farm management practices, lack of regular surveillance, and lack of strict prevention, and control policies. While a meta-analysis of published literature on bTB reported prevalence rates of 2-50% in cattle in India, the true incidence of bTB in India remains ambiguous in the absence of routine national bTB surveillance [8]. Seminal findings in the past showed lower incidences of TB in the indigenous Indian zebu cattle (*Bos indicus*) compared to exotic European cattle (*Bos taurus*) [9-14]. Susceptibility to TB has also been estimated to be influenced by various factors such as herd size, nutritional requirement, age, sex, and dairy farm management practices [9, 15, 16]. A recent study reported significantly higher bTB prevalence in exotic and crossbred cattle than in indigenous cattle breeds [17]. A plethora of studies in the mouse model as well as in humans has indicated that the genetic makeup of a host substantially influences the intracellular survival of mycobacteria by inducing differential immune responses [18-23]. However, a systematic study to compare the underlying immune responses amongst indigenous Indian cattle, and crossbred cattle has not been reported. We hypothesized that higher incidences of bTB in the crossbred cattle might be arising due to inadequate immune response to TB infection compared to the native cattle breeds. To compare the innate cellular responses, first, we segregated the healthy, single intradermal tuberculin test (SITT) negative, and myco-PCR [PCR targeting *M. tuberculosis, M. bovis, M. orygis*, Mycobacterium tuberculosis complex (MTC) and Non-Tuberculous Mycobacteria (NTM) negative cattle from two prominent breeds-indigenous breed Sahiwal, and crossbred-Sahiwal x Holstein Friesian (SHF). Subsequently, we performed a comparative mycobacterial growth assay in the PBMCs isolated from these healthy mycobacterium-naive cattle. Concurrently, we compared the innate immune cytokine responses induced by the PBMCs upon mycobacterial infection and antigenic stimulation. We identified considerable differences in key pro-inflammatory cytokine responses between these breeds that potentially contribute to the differential susceptibility to mycobacterial infection and varied incidence of bTB in these breeds of cattle in India.

## Results

### Higher incidence of tuberculin reactors and myco-PCR positivity in crossbred cattle

To identify, and segregate bTB-negative animals we performed standard SITT, and myco-PCR-based screening of both indigenous Sahiwal breed, and SHF crossbred cattle from a dairy herd **(Fig. 1**). Comparison of SITT response to bovine tuberculin was performed on 24 Sahiwal, and 26 SHF cattle. A total of 9 animals were found to be bovine tuberculin reactors equating to SITT positivity of 18% (9/50). Estimation of the breed-specific tuberculin reactors revealed 4 % (2/50) positivity among Sahiwal cattle, whereas 14 % (7/50) positivity in the case of crossbred SHF cattle (**Fig. 1A**). Concurrently, our myco-PCR methodology that involves a combination of singlet PCRs using previously published primers that detect *M. bovis, M. tuberculosis, M. orygis, MTC*, and pan non-tuberculous mycobacterial (NTM) DNA including the *Mycobacterium avium* complex (MAC) (**Supporting Fig. S1**), and an in-house assembled multiplex-PCR that simultaneously detects and differentiate *M. bovis, M. bovis* BCG, *M. tuberculosis* and pan NTM DNA in the cattle milk and urine samples (**Fig. 1B**) revealed presences of 4% of *M. bovis* positivity (2/50, RD1+, RD4+), 6% *M. tuberculosis* positivity (3/50, RD1+ only), 14% MTC positivity (7/50), and 32% NTM positivity (16/50) (**Table-1**) [24-27]. Altogether, a considerably higher number of crossbred cattle was found to be myco-PCR positive (13/26) compared to the native Sahiwal breed (2/24). **Supporting Table-S1** depicts the detailed distribution of SITT and myco-PCR assay results among all the animals. The primers targeting different mycobacteria are listed in **Supporting Table S2**. We have excluded animals showing SITT positivity, or PCR positivity to either of the mycobacterial species screened in this study for the subsequent evaluation of mycobacterial growth, host cellular responses to mycobacterial infection, and antigenic stimulation (**Table-1 and Supporting Table-S1**). These SITT-negative and myco-PCR-negative cattle were considered mycobacteria-naïve animals that are expected to show primary immune responses when exposed to mycobacterial infection of antigenic stimulation.

**Fig. 1.**
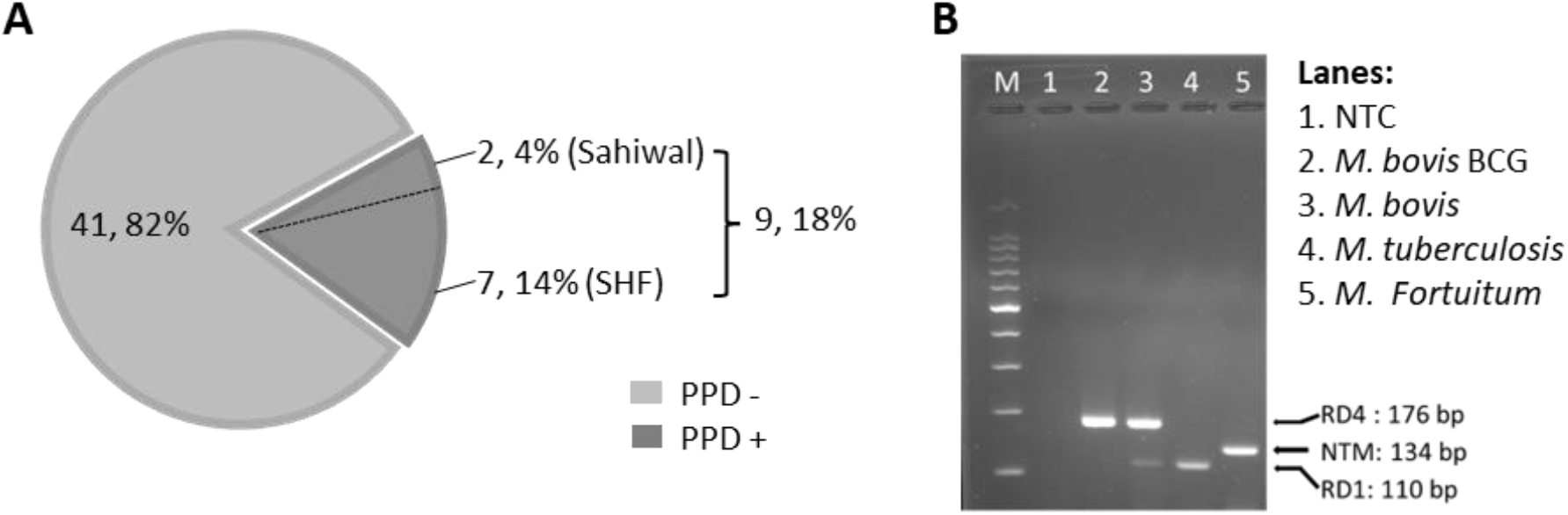
Single Intradermal Tuberculin Test, and PCR-based detection of mycobacterial DNA. (A) The pie chart depicts the incidence of SITT positivity in cattle. A total of 50 cattle were analyzed by SITT. Out of 50 cattle, 9 (18%) were tuberculin reactors. Out of 9 SITT+ cattle, 7 cattle (14%) belonged to Cross Breed and the remaining 2 (4%) were Sahiwal cattle. (B) Agarose gel electrophoresis image of the multiplex PCR using a combination of, - a primer pair targeting specific DNA sequence of the Region of Difference-1 (RD-1) that detects both *M. bovis*, and *M. tuberculosis* but not BCG, or NTMs, - a primer pair targeting the upstream and downstream sequences of the RD-4 region specifically detecting BCG, and *M. bovis* DNA but not *M. tuberculosis* or NTMs, and – a primer pair that detects the presence of pan NTMs DNA.

**Table-1.**
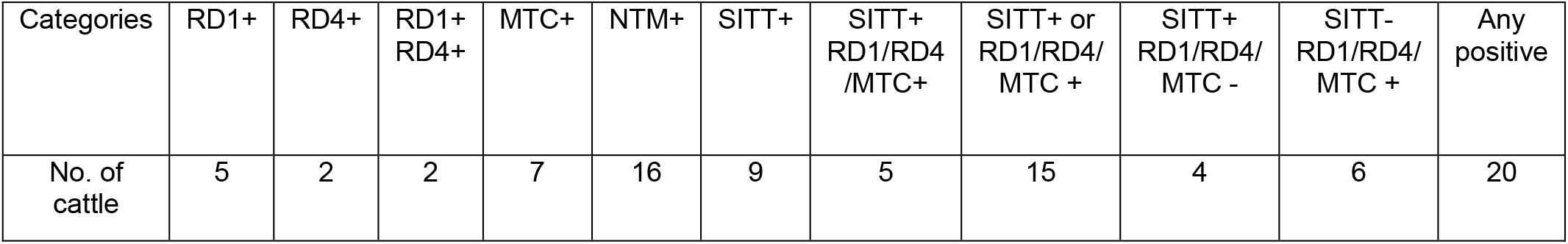
Distribution of SITT and myco-PCR positivity across all the cattle.

| Categories | RD1+ | RD4+ | RD1+<br>RD4+ | MTC+ | NTM+ | SITT+ | SITT+<br>RD1/RD4<br>/MTC+ | SITT+ or<br>RD1/RD4/<br>MTC + | SITT+<br>RD1/RD4/<br>MTC - | SITT-<br>RD1/RD4/<br>MTC + | Any<br>positive |
| --- | --- | --- | --- | --- | --- | --- | --- | --- | --- | --- | --- |
| No. of<br>cattle | 5 | 2 | 2 | 7 | 16 | 9 | 5 | 15 | 4 | 6 | 20 |

### Crossbred cattle PBMCs are conducive to mycobacterial replication

For a comparative evaluation of the permissiveness of the two breeds of cattle to mycobacterial infection, a bovine PBMC-mycobacteria *in vitro* infection assay was established. First, we generated reporter strains of *M. bovis* BCG, and *M. tuberculosis H37Ra* expressing mCherry and tdTomato *via* episomal plasmids *pMSP12::mCherry* and *pTEC27-Hyg*, respectively (**Supporting Table-S3**) [28]. The correlation of the fluorescence of the reporter mycobacterial strains to the CFU was evaluated both in the 7H9 broth culture (**Supporting Fig. S2 A-B**), as well as in the bovine macrophage cells (BOMAC) (**Supporting Fig. S2 C-D**) [29, 30]. The association of the reporter mycobacterial number to total fluorescence was found to be in strong agreement, and a bacterial number-dependent increase in the total fluorescence was observed over five days in both broth culture and BOMAC cell culture (**Supporting Fig. S2)**. A pre-calibrated MOI of 1:10 (Cell: Bacteria) was used for a 5 days-long bovine PBMC culture along with fluorescence measurement at an interval of 24 hours following infection to evaluate the comparative mycobacterial growth in two breeds of cattle (**Fig. 2**). The mean fluorescence of *M. bovis BCG* in the SHF derived PBMC exhibited increasing trend compared to the PBMCs derived from Sahiwal breed of cattle, and at day-5 post-infection the bacterial fluorescence was found to be significantly higher in the former group compared to the later (**Fig. 2A**). Further, PBMCs from Sahiwal breed showed a considerably lower fluorescence for *M. tuberculosis* H37Ra over the course of infection which is significantly lower at day-4 and day-5 post-infection highlighting restricted growth of the bacteria in comparison to the PBMCs from the crossbred SHF cattle (**Fig. 2B**). These observations clearly indicate that indigenous Sahiwal breed of cattle possess significantly greater control over the growth of *M. bovis*, and *M. tuberculosis* strains in comparison to the crossbred SHF cattle.

**Fig. 2.**
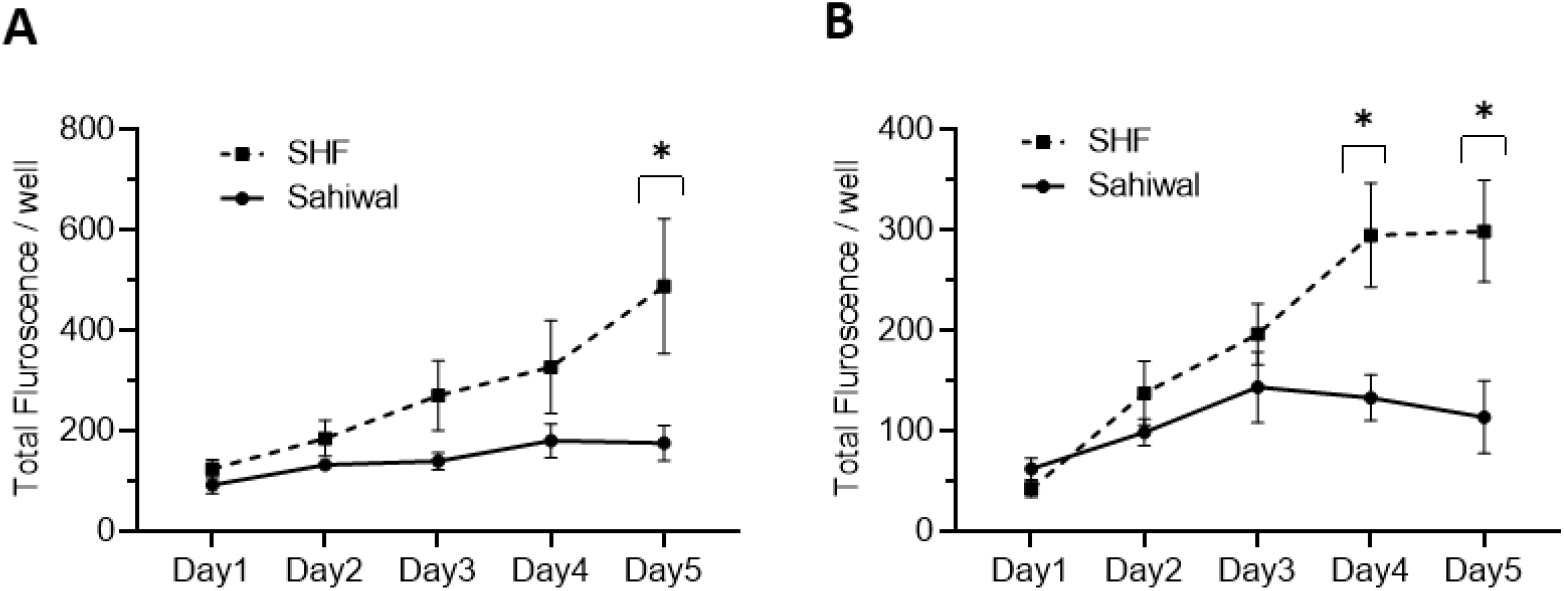
Growth of mycobacteria in bovine PBMC. PBMC from Sahiwal and SHF cattle were infected with fluorescence reporter (A) *M. bovis;* λ_ex_:587nm and λ_em_:610nm, and (B) *M. tuberculosis H37Ra-pTEC27;* λ_ex_:554nm and λ_em_:583nm at an MOI of 1:10 (cell: bacteria). The fluorescence intensity of reporter mycobacteria strains was monitored every 24 hours for 5 days post-infection, and data is depicted as line plots by adjoining the mean ± SEM fluorescence / well measured every day for each mycobacterial strain. n = 6, *,p<0.05 (t-test). The data is representative of two experiments.

### Higher IFN-*γ* production by PBMCs from indigenous cattle breed upon mycobacterial infection, and antigenic stimulation

IFN-*γ* is an important cytokine that regulates innate as well as acquired cell-mediated and humoral immunity to infection by eliciting a number of biological responses in several cell types [31, 32]. IFN-*γ* plays a pivotal role in exerting the host’s protective immunity against Mycobacterial infection [33, 34]. Evaluation of the protein level of IFN-*γ* by ELISA in the bPBMC culture media at 24-hour post-infection with *M. bovis BCG, and M. tuberculosis H37Ra* revealed a significant difference between Sahiwal and SHF cattle (**Fig. 3**). In the first set of experiments, PBMCs from Sahiwal cattle showed a higher production of IFN-*γ* than SHF cattle when exposed to *M. bovis BCG, and M. tuberculosis H37Ra* infection (**Fig. 3A**), while LPS stimulation resulted in a comparable level of IFN-*γ* production by PBMCs from both the sources. This observation was reconfirmed by subsequent experiments wherein in addition to *M. bovis BCG, M. tuberculosis H37Ra* infection, bPBMC were also stimulated with bovine PPD (PPD-B), avium PPD (PPD-A), *M. tuberculosis-* whole cell lysate (WCL), cell wall (CW), and lipoarabinomannan (LAM). We observed that the IFN-*γ* levels at 24-hour post-infection were significantly higher in PBMCs from Sahiwal cattle than SHF cattle in the case of *M. bovis BCG*, and *M. tuberculosis H37Ra* infection, and PPD-B stimulation (**Fig. 3B**). For the rest of the stimulant groups no considerable difference was observed. These observations indicate higher induction of IFN-*γ* by PBMC from indigenous Sahiwal cattle during infection might contribute to the restriction of mycobacterial growth and the resultant lower incidence of TB in this breed of cattle.

**Fig. 3.**
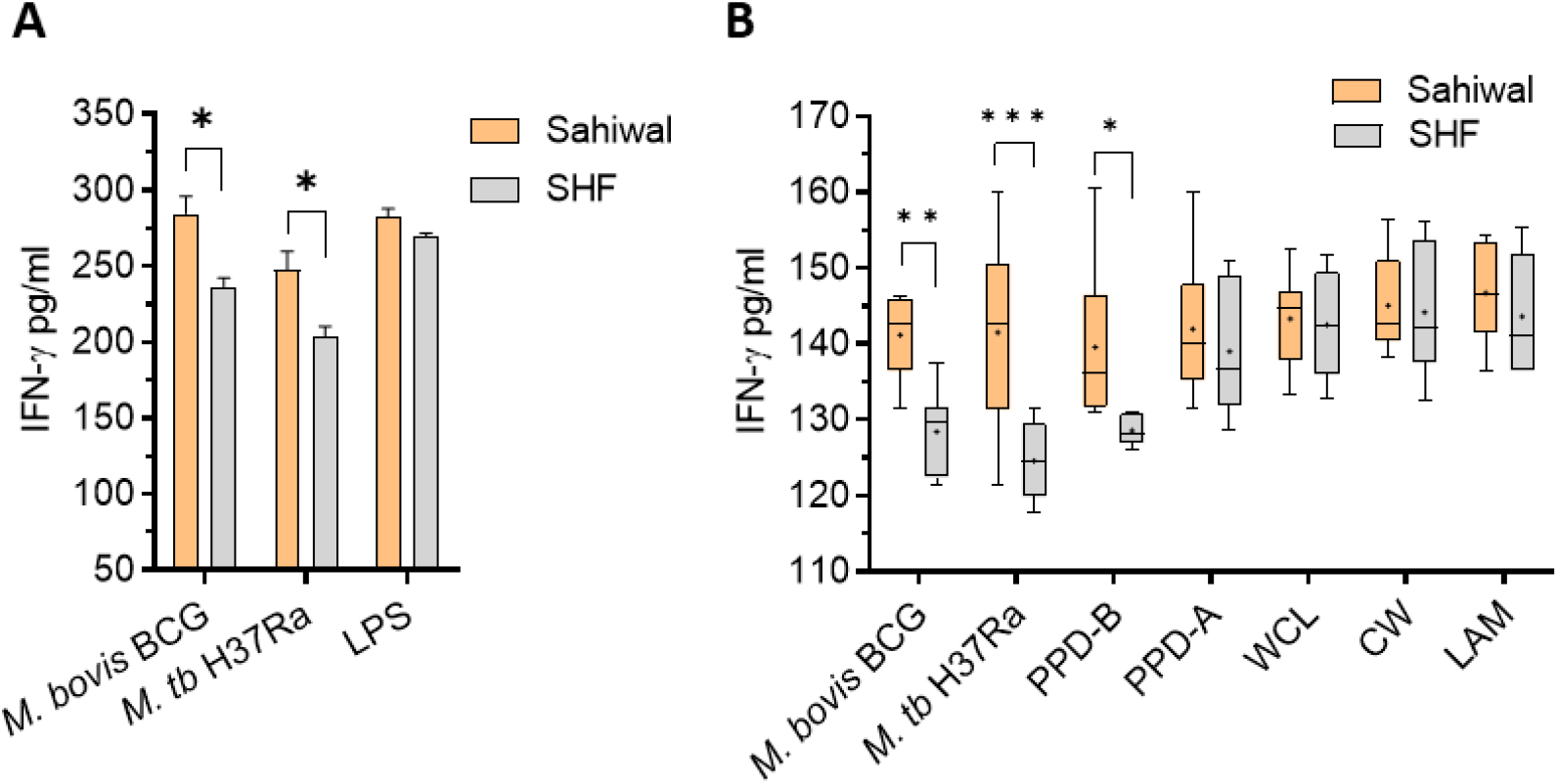
IFN-*γ* response of bovine PBMC to mycobacterial infection and antigenic stimulation. Two sets of experiments were performed. (A) The PBMCs obtained from SITT and myco-PCR negative Sahiwal and SHF cattle were infected with M. bovis BCG or *M. tuberculosis* H37Ra at an MOI of 1:10 (cell: bacteria) or stimulated with Lipopolysaccharide (LPS, 5ug/ml). The bar graph represents IFN-*γ* level in the culture supernatant (pg/ml). Data is mean ± SEM, n=3 animals per group, *,p<0.05. (B) PBMC were either infected with M. bovis BCG or M. tuberculosis H37Ra at an MOI of 1:10 (cell: bacteria) or stimulated with bovine PPD (PPD-B, 300 IU/ml), Avian PPD (PPD-A, 250 IU/ml), M. tuberculosis whole cell lysate (WCL, 5 ug/ml), cell wall (CW, 5 ug/ml) and lipoarabinomannan (LAM, 5ug/ml). IFN-*γ* level was measured at 24 hours post-infection and graphically represented by a box plot, wherein median values are denoted by the horizontal line, the mean is represented by ‘+’, the interquartile range by boxes, and the maximum and minimum values by whiskers. n=6 animals per group. *, P < 0.05; **, P < 0.01; ***, P < 0.001 (t-test). The data is representative of two experiments.

### Transcriptional induction of pro-inflammatory immuno-signature by indigenous cattle PBMCs

Pathogenesis of pulmonary TB largely depends on the orchestration of the players of the cellular immune system and a synchronized interaction of various pro- and anti-inflammatory cytokines at the site of infection [18, 31, 33]. A fine-tuning of multiple cytokines is essential to an effective clearance of the pathogen [35-37]. RNA extracted from the PBMCs from the above experiment at 24 hours post-infection were analyzed for several major cytokines, and signaling molecules by real-time RT-PCR using gene-specific primers (**Supporting Table S4**). These includes IFN-*γ*, IL-17, TNF-α, IFN-β, IL-1β, IL-6, IL-8, cGAS, STING, TBK1, IRF3, and IRF7. **Figure 4** depicts the relative expression of relevant immune-response genes in bar diagrams, and a corresponding heat map showing individual 2^-ddCT^ values for each sample. Of these various immunological mediators, significantly higher transcriptional induction of IFN-*γ* was observed in case of PBMCs from Sahiwal cattle than crossbred cattle when infected with *M. bovis BCG* and *M. tuberculosis H37Ra*, (**Fig. 4A**). In addition, a similar pattern of significantly higher induction was observed in case of IL-17 gene expression by PBMCs from Sahiwal cattle compared to the PBMCs from crossbred SHF cattle when exposed to *M. bovis BCG* and *M. tuberculosis H37Ra* infection (**Fig. 4B**). The serine/threonine kinase TBK-1, which is known for its involvement in the innate immune response to infection by mediating the cGAS-STING-IFN-β axis of cytosolic surveillance pathway, was also found to be significantly upregulated following *M. bovis* BCG infection of the PBMCs from Sahiwal cattle in comparison to the PBMCs from crossbred cattle (**Fig. 4C**). In contrast, IL-1β expression was significantly higher in crossbred cattle PBMCs upon infection with both *M. bovis BCG* and *M. tuberculosis H37Ra*. (**Fig. 4D**). Rest of the genes analyzed in this study exhibited a comparable expression pattern in the case of both breeds of cattle. Stimulation with PPD-B, PPD-A, WCL, CW, and LAM did not exhibit a considerable difference in gene expression between the two breeds of cattle (data not shown). Our findings from quantitative real-time PCR indicates differential transcriptional regulation of important cytokines and signaling pathway such as IFN-*γ*, IL-17, IL-1β, and TBK-1 upon mycobacterial infection may contribute to the differential permissiveness of the two breeds of cattle to TB infection.

**Fig. 4.**
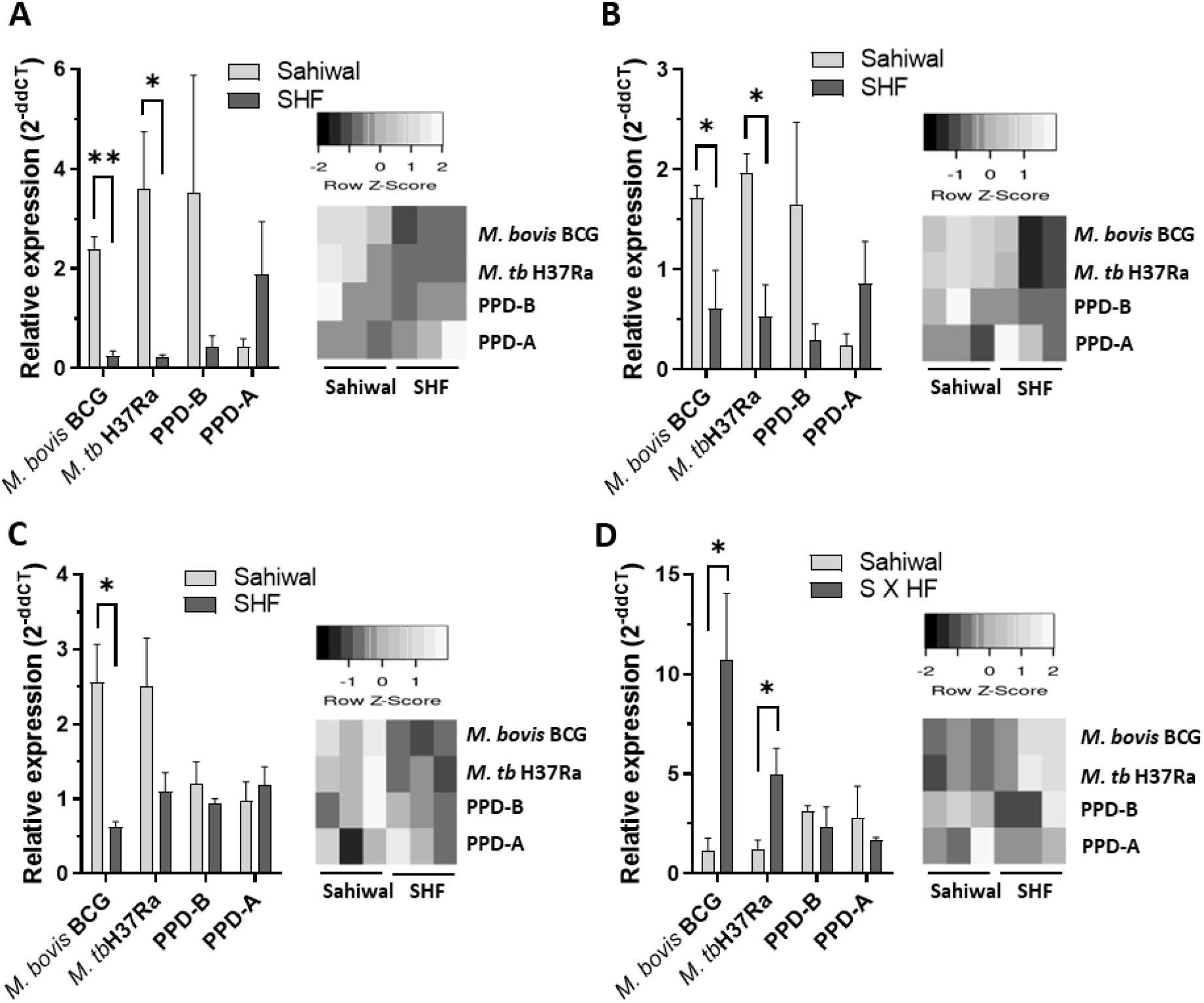
Modulation of host gene expression in bovine PBMC by mycobacterial infection and antigenic stimulation. Expression of various cytokines and immunity-related genes were measured on the RNA extracted from PBMC infected with M. bovis BCG or M. tuberculosis H37Ra at an MOI of 1:10 (cell: bacteria) or stimulated with bovine PPD (PPD-B, 300 IU/ml) and Avium PPD (PPD-A, 250 IU/ml at 24-hour post-infection by semi-quantitative real-time RT-PCR using gene-specific primers. (A) IFN-*γ*, (B) IL-17, (C) TBK-1 and (D) IL-1β. The data were normalized to RPLP0 expression levels and then normalized to the values of uninfected/unstimulated cells to obtain ddCT values. Data is represented as a bar diagram of mean ± SEM of 2-ddCT values as relative expression, n=3, *, p<0.05 (t-test). Each 2-ddCT value was also represented for genes as a heatmap in the inset diagram. Heatmaps were generated using ‘Heatmapper’ software (http://www.heatmapper.ca/expression/). The data are representative of two experiments.

## Discussion

The susceptibility and/or resistance of a host to TB is influenced by multiple factors which include: the nutritional status of the host, age, sex, underlying diseases, host genetic traits, and interaction between the host and the environment [16]. Numerous studies have indicated that genetic diversity among organisms contributes immensely to the differential immune response [21, 22]. A number of studies reported that bovine TB was more prevalent in crossbred cattle compared to the indigenous cattle breeds in India [17, 38]. Thakur and colleagues investigated the prevalence of TB in an organized farm, and a cow shelter in northern India and reported higher TB positivity in Jersey crossbreds [38]. Das and colleagues reported markedly higher incidences of TB in exotic and crossbred cattle (34.6%) compared to indigenous cattle (10.5%) in India [17]. A higher incidence and severity of pathology of bTB in the Holsteins breed compared to zebu breeds was reported in a study conducted in central Ethiopia [10]. As it is apparent that indigenous breeds of cattle have a markedly lower incidence of TB compared to the exotic and crossbred cattle, a comparison of immune responses to TB infection in these cattle may discern the clue of protective immune signature to TB in cattle.

India is home to the largest cattle population in the world with an array of indigenous, and crossbred varieties with enormous genetic variability. Cross-breeding practices remained a preferred approach to enhance the milk yield of indigenous dairy breeds of cattle for more than half a century in India [39]. Especially, the use of European donor breeds such as Holstein Friesian, Jersey, Brown Swiss, Red Dane, etc. for improving non-descript Indian cattle as well as pure-breed indigenous cattle, and the impact of cross-breeding on milk production, reproductive performance, and sustainability in Indian agro-climatic conditions were thoroughly studied via a number of cattle development programs. Exotic inheritance of 1/2 and 5/8 was found to be superior in milk production and sustainability parameters in the majority of the studies compared to other genetic grades [39, 40]. Lower or higher exotic inheritance than the above-mentioned grades did not result in any economic benefit from such cross-breeding practices but rather resulted in unsustainable breed quality in the Indian agro-climatic conditions. As per the last livestock census, crossbred cattle represent more than one-third of India’s total cattle population and contribute to nearly 48% of total cow milk [41]. However, how cross-breeding has influenced the susceptibility and/or resistance trait to different infectious diseases, and the underlying genetic and immunological mechanisms are rarely evaluated systematically.

Here, we studied two of the most prominent dairy breeds of cattle in India, indigenous Sahiwal and crossbred SHF. The crossbred SHF animals included in this study possess 50%-62.5% exotic inheritance. Using both standard SITT assay and myco-PCR, we found a significantly higher incidence of tuberculin reactors-cum-PCR positivity in SHF cattle compared to Sahiwal cattle (**Fig. 1, Table-1**). Our findings are in accordance with the previous studies where a higher percentage of tuberculin reactors was seen in exotic/crossbred cattle than in native cattle [10, 15, 42, 43]. Further, the myco-PCR assays enabled us to detect the presence of MTC and NTM genomic DNA in the milk and urine samples isolated from cattle. The presence of NTM may interfere not only with the SITT readout but also may affect the cellular immune responses of PBMCs isolated from the cattle. Application of such PCR assays provides a cost-effective method to detect not only the species of the infecting mycobacteria in the clinical settings but also allowed us to identify mycobacterial infection-free cattle to be included in the subsequent *ex vivo* PBMC-based experiments.

*Mycobacterium tuberculosis* ensures its survival in the host by replicating primarily in the non-activated macrophages where it inhibits certain essential host processes including the fusion of phagosomes with lysosomes, avoiding its exposure to acidic pH and reactive nitrogen intermediates (RNIs) which are generated for its destruction [18, 31]. Moreover, *M. tuberculosis* blocks antigen presentation, apoptosis, and the suppression of bactericidal activity which occurs as a result of the activation of mitogen-activated protein kinases, IFN-*γ*, and calcium (Ca2+) signaling pathways [44]. Following Toll-like Receptor (TLR-1/2/6) mediated recognition, and phagocytosis of the bacteria, macrophages along with dendritic cells, secrete IL-12 and IL-23, which further stimulates T-cell, and Natural Killer cells to release IFN-*γ* thereby mobilizing the T-helper type 1 (Th1) cytokine pathway. IFN-*γ* is the key cytokine indispensable in defense against TB. IFN-*γ* activated macrophages enhance the microbicidal activity of macrophages by allowing the formation of phagolysosomes wherein the mycobacteria are deprived of essential nutrients such as iron, and exposed to anti-microbial peptides, and reactive oxygen or nitrogen intermediates [32, 45, 46]. Lower production of IFN-*γ* indicates a reduced activity of macrophages which promotes mycobacterial growth [32, 37]. The findings from ELISA and real-time RT-PCR indicated a significantly higher induction of IFN-*γ* response by PBMCs from Sahiwal cattle compared to crossbreed cattle PBMCs indicating induction of superior anti-TB immune responses in the native Sahiwal cattle (**Fig. 4**). This observation was supported by the lower growth of both *M. bovis BCG* and *M. tuberculosis H37Ra* strains in the PBMC cultures of Sahiwal cattle compared to the crossbred SHF cattle (**Fig. 2**).

Although IFN-*γ* plays a key role in the defense against TB, this cytokine alone can’t generate the necessary immune response to provide protection against TB. The TB disease progression is controlled by a coordinated network of several different cytokines, chemokines, and signaling molecules. We analyzed the mRNA levels of a number of pro-inflammatory, anti-inflammatory, and immuno-regulatory mediators in addition to IFN-*γ* such as IL-17, TNF-α, IFN-β, IL-1β, IL-6, IL-8, cGAS, STING, TBK1, IRF3, and IRF7. While IL-17, IL-1 β and TBK1 exhibited considerable differential regulation in the mRNA levels, a comparable level of expression was observed for the rest of the immune mediators (**Fig. 4**). A significantly higher induction of IL-17 was exhibited by PBMCs from Sahiwal cattle compared to crossbreed cattle PBMCs (**Fig. 4b**). IL-17 is another key cytokine involved in exhibiting protective immunity against *M. tuberculosis* infection. It plays a major role in combating the growth of the tubercle bacilli by promoting a Th1-biased immune response [47]. The differentiation of Th17 cells occurs as a result of an increase in the level of pro-inflammatory cytokines such as IL-6, IL-21, IL-1beta, and TNF-a [36, 47-49]. IL-17 induces the recruitment of neutrophils, macrophages, and Th1 cells to the site of inflammation. IL-17 also restricts the growth of Mycobacteria by inducing the expression of various chemokines, and by recruiting IFN-*γ*-producing cells [35]. In contrast, IL-1β expression was significantly higher in crossbred cattle PBMCs upon mycobacterial infection (**Fig. 4c, 4d**). While IFN-*γ* and IL-17 primarily promote macrophage activation, granuloma formation, and clearance of intracellular tubercle bacilli, IL-1β has been implicated in aggravated inflammation [50]. Further, phosphorylation of TBK-1 is involved in a plethora of intracellular signaling events including the cGAS-STING-IFN-β axis of the cytosolic surveillance pathway of the host to respond to invading infectious agents including regulation of cell proliferation, autophagy, and apoptosis [51, 52]. While the role of TBK-1 in antiviral response is well documented, divergent views on the anti-bacterial effect especially anti-mycobacterial responses are linked to this cytosolic kinase [53-56]. Several studies reported the essentiality of the TBK-1 phosphorylation-mediated activation of the IRF-1 pathway for mycobacterial clearance, while others associated it with higher immunopathology [54, 55]. These observations indicate the necessity of tightly regulated TBK-1-mediated signaling for a host-favored immune response.

In some, our study elucidated the association of important mediators of immune responses with the differential TB susceptibility phenotype of the indigenous Sahiwal cattle and the crossbred SHF cattle by employing *ex vivo* bovine PBMC-mycobacterial infection model using *M. bovis* BCG vaccine strain, and *M. tuberculosis* H37Ra strain. Especially, our study highlighted that divergence in the expression of host factors such as IFN-*γ*, IL-17, IL-1β and TBK-1 potentially play a major role in determining the degree of susceptibility to mycobacterial infection in cattle. However, further studies with virulent human and bovine tubercle bacilli (in this study BSL2 grade mycobacteria-*M. bovis BCG* and *M. tuberculosis H37Ra* have been used), a greater number of indigenous and crossbred cattle, and application of genomic, transcriptomic and proteomic approaches would elucidate the association of the global immune response signature to TB susceptibility and/or resistance in cattle. Our study also highlighted the importance of the use of PCR-based mycobacterial DNA detection and differentiation of mycobacteria species in studies that allow the identification of mycobacterial infection-naïve cattle to evaluate the innate immune response of PBMCs to mycobacterial stimulation or infection. Further, a highly sensitive multiplex-PCR-based assay would be immensely useful for screening cattle herds as well as human clinical TB cases that would aid in the epidemiological characterization of causative mycobacterial species and devising appropriate treatment strategies.

Finally, this study is an important step forward toward identifying the association of TB susceptibility to the underlying innate immune responses in indigenous and crossbred cattle in India. This study addresses an extremely important yet untouched aspect of bTB scenario in Indian cattle which is identifying host factors conferring susceptibility and/or resistance to TB that remained one of the biggest public health problems for centuries. Our results not only provide proof-of-concept data for the hypothesis that genetic variability of bovine due to breed variation influences TB susceptibility and resistance but also provide a reason for adopting an appropriate crossbreeding policy that balances production and disease resistance traits for sustainable livestock farming.

## MATERIALS AND METHODS

### Ethics statement

All the experiments were reviewed and approved by the Institutional Biological Safety Committee (IBSC, Approval No. IBSC/2018/NIAB/BD/001) of the National Institute of Animal Biotechnology, Hyderabad, and animal experiments were approved by the Animal Ethics Committee of the West Bengal University of Animal and Fishery Sciences, Kolkata, India [Approval No. IAEC/22 (B), CPCSEA Reg. No.763/GO/Re/SL/03/CPCSEA), Committee for the Purpose of Control and Supervision on Experiments on Animals, India]. Blood collection from the jugular vein and tuberculin test on cattle were performed by a trained Veterinarian, and all the procedures were performed following relevant guidelines and regulations.

### Bacteria, Plasmids, and generation of reporter Mycobacterial strains

The mycobacterial strains and plasmids used in the study are listed in **Supplementary Table -S1**. *M. bovis* BCG (Danish 1331 sub-strain) was procured from NIBSC, UK, and *M. tuberculosis* H37Ra strain was kindly provided by Prof. S. Banerjee, University of Hyderabad, India. Non-Tuberculous Mycobacteria (NTM) *M. fortuitum* was procured form MTCC, CSIR-IMTECH, Chandigarh, India. Mycobacterial strains were grown to mid-log phase in Middle Brook (MB) 7H9 media and glycerol stocks were prepared and stored at -80°C as described earlier [57]. For *in vitro* infection, fresh bacterial cultures were grown to the mid-log phase, and bacterial cells were washed thoroughly with 1XPBS, and finally resuspended in cell-culture media following pre-calibrated dilutions, as described previously [54]. For generating Mycobacterial reporter strains first, electrocompetent cells of *M. bovis* BCG and *M. tuberculosis* were prepared as described previously [54], and transformed with pMSP12::mCherry (was a gift from Lalita Ramakrishnan, Addgene plasmid # 30169; http://n2t.net/addgene:30169), and pTEC27-Hyg (was a gift from Lalita Ramakrishnan, Addgene plasmid # 30182; http://n2t.net/addgene:30182) [28]. Briefly, 100ul of competent Mycobacterial cells were mixed with 100ng of plasmid DNA, and transferred to a Micro Pulser Electroporation Cuvettes cuvette (Biorad # 1652086 with 0.2 cm gap) and pulsed at 2500V, 25uF capacitance, and 1000 Ω resistance using an electroporator (Gene Pulser Xcell Microbial System #1652662, Bio-Rad). The cells were immediately aspirated and inoculated in 2ml of MB-7H9 broth without any selective antibiotics and incubated at 37°Cunder shaking conditions for 48 hours. Subsequently, bacterial cells are plated onto MB-7H11 agar containing selective antibiotics-25 µg/ml of Kanamycin, and 150 µg/ml of Hygromycin, respectively, and incubated at 37°C for 4 weeks. Transformed bacterial clones grown on the selective plates were detected by colored colonies, and also confirmed for the presence of plasmid by colony PCR, and subsequently confirmed by fluorescence microscopy. Glycerol stocks of the reporter mycobacterial strains were prepared, and stored for future use. All the mycobacterial strains used in this study were of the BSL-2 category, and these are cultured in a BSL-2 laboratory.

### Genomic DNA extraction from mycobacteria

Standard methodology was followed for genomic DNA extraction from various mycobacteria as described previously [58]. Briefly, the bacterial culture was incubated with 1% glycine in a 37°Cshaker for 24 hours. After incubation, the cells were harvested by centrifuging the bacteria at 8000 rpm for 10 minutes. The cells were resuspended in 5ml of TEG (Tris 25mM pH 8.0, EDTA 10mM pH 8.0, Glucose 50 mM) solution and mixed gently. 500ul of lysozyme (10mg/ml) was added to this suspension and incubated overnight at 37°Cshaker incubators. Later, 1 ml of 10% SDS and 500ul of Proteinase K (20mg/ml) were added to the cell lysate and incubated at 55°Cfor 40 minutes. Subsequently, a solution comprising 2ml of 5M NaCl and 1.6ml of pre-warmed 10% CTAB was added to the cell lysate which was later incubated at 65°Cfor 10 minutes. The suspension was then centrifuged at 12000 rpm for 30 minutes at room temperature. The supernatant was subjected to phenol-chloroform extraction twice, and genomic DNA was precipitated by the addition of isopropanol. The DNA pellet was washed twice with 70% ethanol, air dried, and resuspended in autoclaved distilled water and stored at -20°C. The genomic DNA was used as a template for PCR experiments. Genomic DNA from *Mycobacterium bovis* BCG, *M. tuberculosis* H37Ra, and *M. fortuitum* were extracted by the above method. Genomic DNA extracted from *Mycobacterium bovis* irradiated whole cells (#NR-31210), and purified Genomic DNA of *M. tuberculosis* H37Rv strain (#NR-13648) procured from BEI resources, USA was used as PCR templates.

### DNA extraction from cow milk and urine

Genomic DNA was isolated from cow milk and urine samples as described previously [2]. Briefly, milk samples were centrifuged at 12,000 x g for 20 minutes at room temperature. A sterile cotton swab was used to remove the fat layer, and the supernatant was discarded. The pellet was vortexed and subsequently resuspended in 500ul of IRS (Inhibitory Removing Solution with pH (7.4) containing 25M guanidium isothiocyanate, 0.025M EDTA, 0.05M Tris, 0.5% Sarkosyl and 0.186M β-mercaptoethanol). Later the samples were incubated at 37°Cfor 60 minutes. After incubation, the samples were again centrifuged at 12,000 x g for 10 minutes and the supernatant was discarded. The pellet was washed with water once and resuspended in 100µl of lysis buffer (10% Chelex 100, 0.3% tween 20, 0.03% Triton X100). The samples were then incubated at 90°Cfor 40 minutes and subjected to another round of centrifugation at 10,000 x g for 10 minutes at room temperature. The supernatant was collected and used as template DNA for PCR reactions, or stored at -20°Cfor future use.

### Single and multiplex-PCR assay for mycobacterial DNA detection

PCR assays were carried out targeting the genomic DNA of single mycobacterial species as well as multiple species in a single reaction. We observed higher sensitivity of single species /gene-specific PCR compared to the multiplex-PCR assay. Comparative genomics has revealed that based on the presence or absence of regions of difference (RD) mycobacterial species or strains can be differentiated. In this study, a combination of previously published primers targeting specific sequences of RD1, RD4, RD12, and *rpoB* gene was used to develop the single and multiplex-PCR to detect and differentiate *M. tuberculosis, M. bovis, M. bovis BCG, M. orygis, pan-MTC* and pan-*NTM (*including MACs) in the cattle samples (**Supporting Fig. S1**). **Supporting Table-S1** shows the list of mycobacterial genomic region-specific primers used in this study. The PCR protocol was executed using the Sapphire PCR master mix (TAKARA) following manufacturer protocol. Subsequently, PCR amplicons were analyzed by agarose gel electrophoresis and visualized using a UV trans-illuminator. The expected size of amplicons with RD1, RD4, RD12-*M. orygis*, MTC, and NTM primers are 110bp, 176bp, 264bp, 235bp, and 134bp, respectively (**Supporting Fig. S1)**. While the multiplex-PCR could detect mycobacteria in .1ng of template DNA, the single species PCR could detect mycobacteria in .01ng of total genomic DNA extracted from cattle milk or urine.

### Single Intradermal Tuberculin Test (SITT)

Fifty cows from two adjacent herds were subjected to the single intradermal tuberculin test. The neck region of the animals was shaved and the thickness of the skin was measured with the use of a caliper before injecting bPPD. One hundred microliters (0.1ml) of bovine PPD (2,000 U/animal of bPPD) (obtained from Indian Veterinary Research Institute, Izatnagar, India; 1mg protein/ml) was injected into the skin of the cervical region. Seventy-two-hour post bPPD injection skin induration was evaluated by measuring the skin thickness with a caliper. The result was graded as bPPD positive reactor when differences in the skin thickness at the injection sites are at least 5 mm or greater.

### *In vitro* bovine macrophage cell culture and mycobacterial growth assay

Bovine macrophage cell line-BOMAC was used for calibrating the infection dose (MOI), and evaluating the association of fluorescence measurements of the reporter mycobacterial strains with the number of bacteria over the period of infection [29, 30]. BOMAC cells were maintained in DMEM supplemented with 10% heat-inactivated Fetal Bovine Serum (FBS). BOMAC cells were infected with the reporter strains of *M. tuberculosis H37Ra-pTEC2*, and *M. bovis BCG*-mCherry at different MOIs. The cells were infected for 3 hours, subsequently, the cells were washed thoroughly, and incubated in a TC incubator in fresh media. The fluorescence intensity was measured at λex/ λem 554/581nm (*M. tuberculosis H37Ra-pTEC27*), and 587/610nm (*M. bovis BCG*-mCherry) respectively at an interval of 24 hours daily for 5 days using a fluorescence multimode plate reader (Biorad). An MOI of 1: 10 was considered for *ex vivo* mycobacterial growth in the bPBMC.

### *Ex vivo* bovine PBMC culture and mycobacterial growth assay

Blood samples were collected from clinically healthy, SITT-negative, and myco-PCR-negative cattle. Five ml of blood was collected from the jugular vein of each animal, and bPBMC was isolated using Histopaque®-1077 (Sigma) following manufacturer recommendations. Briefly, blood was diluted with DPBS at a ratio of 1:1. The sample was layered slowly on top of the Ficoll density gradient buffer (5 ml diluted Blood on 3 ml Ficoll) and centrifuged at 400 x g for 30 minutes. The monocyte layer was carefully separated and washed twice with DPBS at 200 x g for 10 minutes. Contaminating RBCs were removed from the cell suspension by adding RBC lysis buffer (Sigma-Aldrich) and following manufacturer protocol. Purified bPBMC were then finally suspended in 5ml of DMEM complete media containing antibiotics, and added to an ultra-low attachment 6-well plate. The cells were incubated in a TC incubator for 24 hours. For mycobacterial growth assay, bPBMC were seeded on to 96-well TC plate at a density of 5×10^4^ cells/well. The cells were infected with mid-log phase cultures of reporter mycobacteria (*M. tuberculosis H37Ra-pTEC27* and *M. bovis BCG-mCherry)* at a pre-calibrated MOI of 1:10 and were incubated in a TC incubator. The fluorescence intensity was measured at λex/ λem 554/581nm (*M. tuberculosis H37Ra-pTEC27*), and 587/610nm (*M. bovis BCG*-mCherry) respectively at an interval of 24 hours daily for 5 days using a fluorescence multimode plate reader.

### Infection and antigenic stimulation of bovine PBMC for evaluation of innate immune responses

For the evaluation of innate immune responses, bPBMC were seeded onto 24-well TC plates at a density of 2×10^5^ cells/well, and infected with either of the mycobacteria (*M. tuberculosis H37Ra, or M. bovis BCG*) at a pre-calibrated MOI of 1:10, or cells were stimulated with LPS (1µg/ml), PPD-B (300 IU/ml), PPD-A (250 IU/ml), *M. tuberculosis* H37Rv Whole Cell Lysate (WCL, 5 µg/ml), cell wall (CW, 5 µg/ml) and purified Lipoarabinomannan (LAM, 5µg/ml). Twenty-four hours post-infection cell culture supernatants were separated for measurement of IFN-*γ* protein levels, and total RNA was extracted from bPBMC for measurement of the mRNA transcripts levels of major cytokines, chemokines, and innate immune-signaling mediators.

### RNA extraction, and real-time RT-PCR

Total RNA was extracted from bPBMC using RNeasy Plus Kit (Qiagen Inc, CA, USA) following manufacturer protocol. Contaminating genomic DNA was removed by additional treatment with RNase-free DNase (Qiagen Inc, CA, USA). The quality and quantity of RNA were analyzed using a NanoDrop Spectrophotometer (Thermo Scientific). cDNA was synthesized from RNA using the Prime script 1^st^-strand cDNA synthesis kit (Takara) as per the manufacturer’s instructions and using a mixture of random hexamer and oligo dT primers. Primers were designed for bovine gene targets (IFN-*γ*, IL-17, TNF-α, IFN-β, IL-1β, IL-6, cGAS, STING, TBK1, IRF3, and IRF7) (**Supporting Table-S4**) using Primer-BLAST (NCBI) and real-time PCR was performed using a CFX96 Touch System (Biorad). Real-time PCR protocol started with an initial denaturation and enzyme activation at 95°Cfor 2 minutes followed by 40 cycles of denaturation at 95°Cfor 15 seconds, annealing and extension were carried out for 1 minute at a temperature ranging from 55°C to 65°C (based on the target gene). Melt curve analysis was performed by heating the samples from 65°C to 95°Cwith an increment of 0.5 and fluorescence was recorded. Relative gene expression of the target genes was calculated using the 2^−ΔΔCT^ method with RPLP0 as an internal control.

### Bovine cytokine Enzyme-Linked Immunosorbent Assay

Twenty-four hours post-infection cell culture supernatants were separated, filtered through 0.2 µ membrane plate filters, and subjected to ELISA for the measurement of IFN-*γ* protein levels using bovine IFN-*γ* specific sandwich ELISA kit as per the manufacturer protocol (K04-0002, Krishgen Biosystems). The absolute concentrations were estimated by referring to a standard curve and expressed as picogram per milliliter.

### Statistical Analysis

GraphPad Prism 9 was used to perform the statistical analysis and preparation of the graphs. For comparison of group means t-tests were performed, and differences were considered significant when p<0.05. All the results are shown as the mean ± SEM, unless otherwise described in the figure legends. Heatmaps of the gene expression data were generated using ‘Heatmapper’ software (http://www.heatmapper.ca/expression/).

## Acknowledgments

Financial support from the NIAB intramural grant, and the Department of Biotechnology (DBT), Govt. of India (Grant No. BT/PR31378/AAQ/1/745/2019) are thankfully acknowledged. Support by DBT for providing Junior Research Fellowship to RK and AR; Department of Science and Technology (DST), Govt. of India for providing the Inspire fellowship (JRF) to SG. Prof. Sharmistha Banerjee, University of Hyderabad, India is thankfully acknowledged for providing *the M. tuberculosis* H37Ra strain.

## Author Contributions

Conceived and designed the experiments: BD. Performed the experiments: RK, SG, AR, HKM, US, BD. Analyzed the data: RK, SG, US, BD. Contributed reagents/ materials/analysis tools/ facility: BD, US. Wrote the paper: RK, SG, US, BD. Provided overall supervision throughout the study: BD.

## Supporting information

Please see the supporting information file.

