## Supplementary information for "Divergent proinflammatory immune responses associated with the differential susceptibility of cattle breeds to Tuberculosis"

### Affiliations:

### *Corresponding Author:

**Supporting Fig. S1**

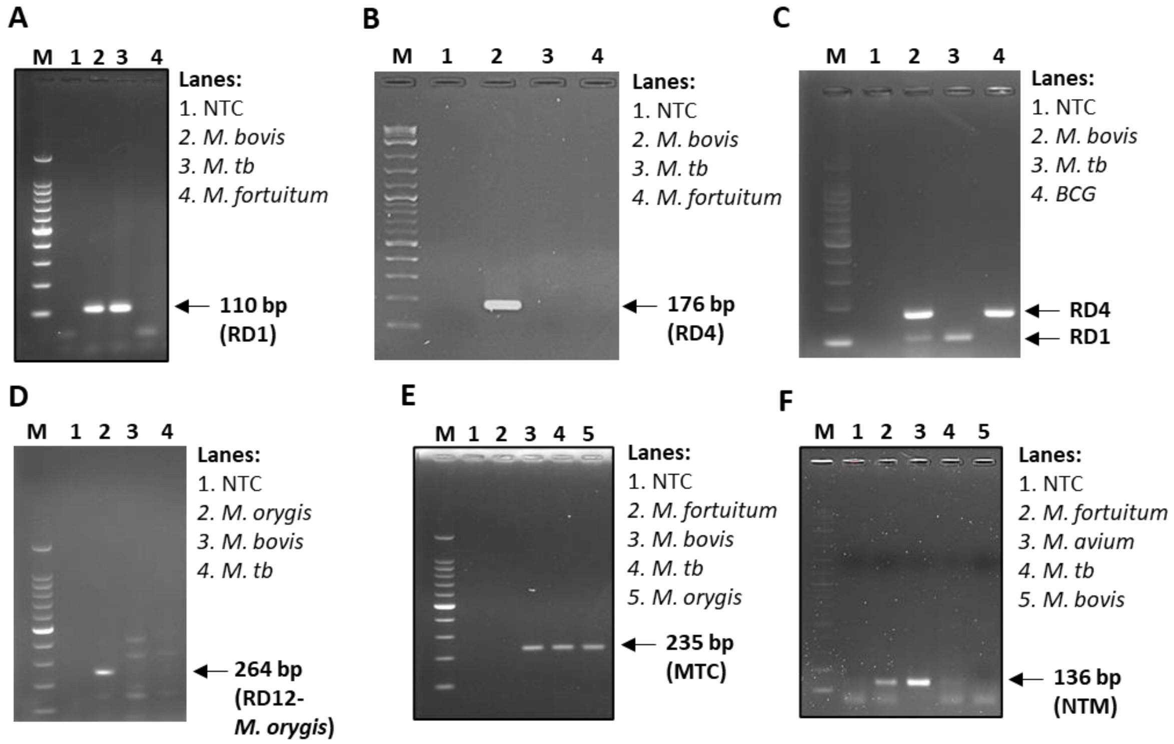


**Supporting Fig. S1. PCR-based detection of mycobacterial DNA.** Representative agarose gel electrophoresis images of the PCR analysis targeting specific genomic regions of mycobacteria to detect (A) RD1 region- positive for *M. tuberculosis* and *M. bovis*, (B) RD4 region- positive of *M. bovis* and BCG, (C) RD1 and RD4 combined region to differentiate *M. tuberculosis*, *M. bovis,* and BCG, (D) RD12- *M. orygis,* (E) pan MTC, and (F) pan- NTM. Genomic DNA from different species of mycobacteria was used as the positive template for PCR. NTC- non-template control.

**Supporting Fig. S2**

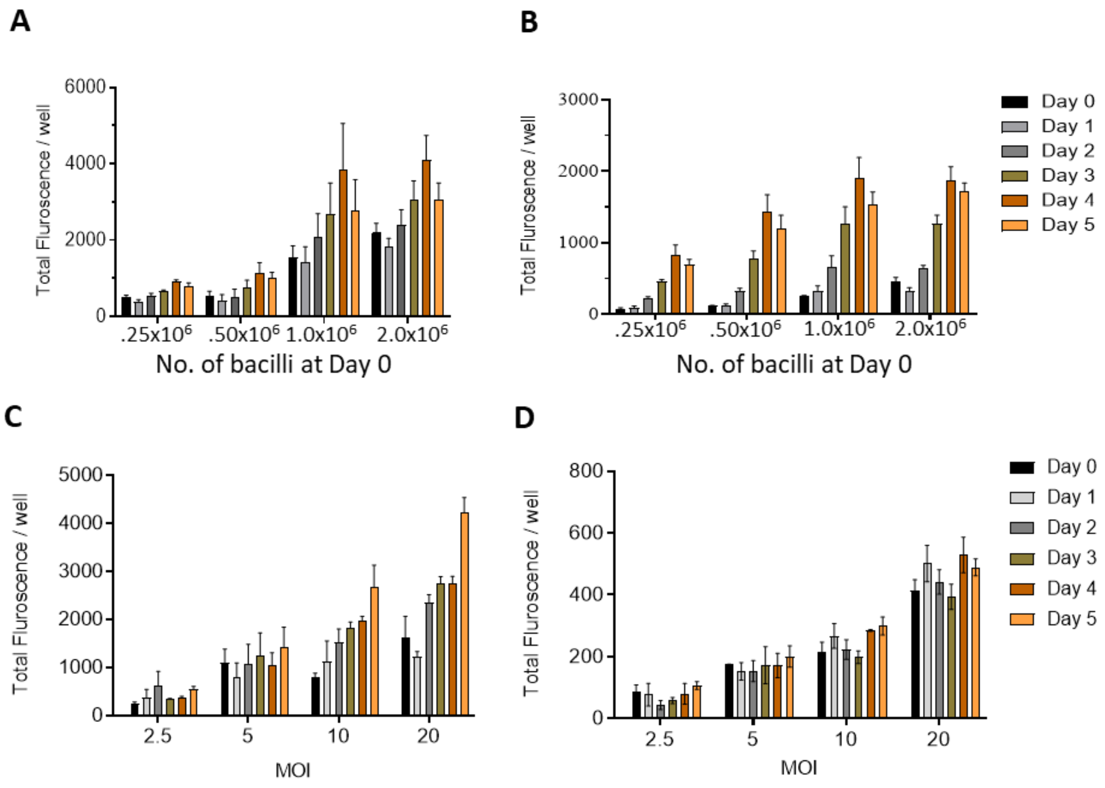


**Supporting Fig. S2. Association of fluorescent reporter mycobacterial number with the total fluorescence.** The association of the fluorescence of the reporter mycobacterial strains- *M. bovis* BCG-mCherry and *M. tuberculosis H37Ra-tdTomato* expressing mCherry and tdTomato to the CFU were evaluated both in the (A, B) 7H9 broth culture and (C, D) BOMAC cells, respectively. The bacterial growth rate in 7H9 broth was determined by plotting the total fluorescence intensity/well against the different numbers of bacteria: 0.25 M, 0.5 M, 1 M, and 2 million at day-0 of the culture, and monitored for 5 days. For growth in the BOMAC cells, cells were infected with the reporter strains at different MOI (2.5, 5, 10, and 20) and the fluorescence intensity was monitored every day for 5 days post-infection. The fluorescence intensity was measured at λex/ λem 554/581nm (*M. tuberculosis H37Ra-pTEC27*), and 587/610nm (*M. bovis BCG*-mCherry). The Pearson's correlation coefficient (*r*) between the bacterial CFU and fluorescence intensity was found to be strongly positive (r = 0.874-0.967, and 0.841-0.990 for *M. tuberculosis H37Ra-tdTomato* in 7H9 broth and BOMAC culture, respectively over 5 days; r = 0.780-0.990, and 0.974-0.999 for *M. bovis BCG-mCherry* in 7H9 broth and BOMAC culture, respectively). An MOI of 1: 10 was considered for *ex vivo* mycobacterial growth in the bPBMC.

| Supporting Table- S1: PCR primers used for mycobacterial detection | | |
|---|---|---|
| **Name** | **Nucleotide Sequence** | **Reference** |
| RD1 | F, 5’-CCCTTTCTCGTGTTTATAGTTTGA-3’<br>R, 5’-GCCATATCGTCCGGAGCTT-3’ | 27 |
| RD4 | F, 5’-AATGGTTTGGTCATGACGCCTTC-3’<br>R, 5’-CCCGTAGCGTTACTGAGAAATTGC-3’ | 26 |
| RD12-<br>*M. orygis* | F, 5’-GTGGAAATGGAAGCGTTGACC-3’<br>R, 5’-GGTACCTCCTCGATGAACCAC-3’ | 25 |
| MTC | F, 5’-CGTACGGTCGGCGAGCTGATCCAA-3’<br>R, 5’-CCACCAGTCGGCGCTTGTGGGTCAA-3’ | 24 |
| NTM | F, 5’-GGAGCGGATGACCACCCAGGACGTC-3’<br>R, 5’-CAGCGGGTTGTTCTGTCCATGAAC-3’ | 24 |

| **Supporting Table- S2: Animal-wise distribution of SITT and PCR positivity.** | | | | | | | | | | |
|---|---|---|---|---|---|---|---|---|---|---|
| **Animal code No** | **RD1+** | **RD4+** | **MTC+** | **NTM +** | **SITT+** | **SITT+ RD1/RD4/ MTC+** | **SITT+ or RD1/RD4/ MTC +** | **SITT+ RD1/RD4/ MTC -** | **SITT- RD1/RD4/ MTC +** | **Any positive** |
| **S1** | | | | | | | | | | |
| **S2** | | | + | + | + | + | + | | | + |
| **S3** | | | | + | + | | + | + | | + |
| **S4** | | | | | | | | | | |
| **S5** | | | | | | | | | | |
| **S6** | | | | | | | | | | |
| **S7** | | | | | | | | | | |
| **S8** | | | | | | | | | | |
| **S9** | | | | | | | | | | |
| **S10** | | | | | | | | | | |
| **S11** | | | | | | | | | | |
| **S12** | | | | | | | | | | |
| **S13** | | | | | | | | | | |
| **S14** | | | | | | | | | | |
| **S15** | | | | | | | | | | |
| **S16** | | | | | | | | | | |
| **S17** | | | | | | | | | | |
| **S18** | | | | | | | | | | |
| **S19** | | | | | | | | | | |
| **S20** | | | | | | | | | | |
| **S21** | | | | | | | | | | |
| **S22** | | | | | | | | | | |
| **S23** | | | | | | | | | | |
| **S24** | | | | | | | | | | |
| **SHF1** | | | + | + | | | + | | + | + |
| **SHF2** | + | + | + | + | | | + | | + | + |
| **SHF3** | | | | + | | | | | | + |
| **SHF4** | | | | + | | | | | | + |
| **SHF5** | + | | + | + | + | + | + | | | + |
| **SHF6** | + | | + | + | | | + | | + | + |
| **SHF7** | | | + | + | | | + | | + | + |
| **SHF8** | | | | + | | | | | | + |
| **SHF9** | + | + | | + | | | + | | + | + |
| **SHF10** | + | | | | | | + | | + | |
| **SHF11** | | | | | | | | | | |
| **SHF12** | | | + | | + | + | + | | | + |
| **SHF13** | | | | | | | | | | |
| **SHF14** | | | | | | | | | | |
| **SHF15** | | | | | | | | | | |
| **SHF16** | | | | | | | | | | |
| **SHF17** | | | | | | | | | | |
| **SHF18** | | | | | + | | + | + | | + |
| **SHF19** | | | | | + | | + | + | | + |
| **SHF20** | | | | | + | | + | + | | + |
| **SHF21** | | | | + | + | + | + | | | + |
| **SHF22** | | | | + | + | + | + | | | + |
| **SHF23** | | | | + | | | | | | + |
| **SHF24** | | | | | | | | | | |
| **SHF25** | | | | + | | | | | | + |
| **SHF26** | | | | + | | | | | | + |
| **Total** | **5** | **2** | **7** | **16** | **9** | **5** | **15** | **4** | **6** | **20** |
| +, positive of the given type; Blank cell, negative for the given type | | | | | | | | | | |

| **Supporting Table- S3: Plasmids, mycobacterial strains, and mycobacterial antigens/cell components.** | | |
|---|---|---|
| **Name** | **Description** | **Source** |
| **Plasmids** | | |
| pMSP12::mCherry | Mycobacterial reporter plasmid expressing mCherry | Addgene plasmid # 30169 |
| pTEC27-Hyg | Mycobacterial reporter plasmid expressing tdTomato | Addgene plasmid # 30182 |
| ***Mycobacterial*** **strains** | | |
| *M. tuberculosis H37Ra* | *M. tuberculosis* H37Ra strain- *BSL2 grade laboratory strain* | *Prof. Sharmistha Banerjee, University of Hyderabad* |
| *M. bovis BCG* | *M. bovis BCG Danish 1331 strain* | NIBSC, UK |
| *M. tuberculosis* H*37Ra*-tdTomato | *M. tuberculosis* H37Ra strain- *harbouring* pTEC27-Hyg | *This study* |
| *M. bovis BCG* - mCherry | *M. bovis BCG Danish 1331* strain harboring pMSP12::mCherry | *This study* |
| *M. fortuitum* | Non-tuberculous mycobacterium strain | *MTCC repository, CSIR-IMTECH, India* |
| **Mycobacterial antigens/cell components** | | |
| *PPD-A* | Purified protein derivative from *M. avium* | *ThermoFisher* |
| *PPD-B* | Purified protein derivative from *M. bovis* | *ThermoFisher* |
| *WCL* | *M. tuberculosis whole cell lysate* | *BEI Resources, USA* |
| *CW* | *M. tuberculosis cell wall* | *BEI Resources, USA* |
| *LAM* | *M. tuberculosis Lipoarabinomannan* | *BEI Resources, USA* |

| Supporting Table- S4: Real-time PCR primers (bovine genes) | | |
|---|---|---|
| **Primer Name** | **Nucleotide Sequence** | **Gene description** |
| *bIFN-γ | F, 5′-gctgattcaaattccggtgga-3′<br>R, 5′-caggcaggaggaccattacg-3′ | Interferon-gamma |
| bIL-17A | F, 5′-agaaggcccaccgattatca-3′<br>R, 5′-ccaccttcccttcagcattga-3′ | Interleukin 17 |
| bIL-1β | F, 5’-cagtgcctacgcacatgtct-3′<br>R, 5′-ccagggatttttgctctctg-3′ | Interleukin 1 beta |
| bIL-6 | F, 5′-cagctatgaactcccgcttc-3′<br>R, 5′-ttcggttttctctggagtgg-3′ | Interleukin 6 |
| bcGAS | F, 5′-ttcaaaggcgtagacctgct-3′<br>ccgataagattcccccttct-3′ | Cyclic GAMP synthase |
| bSTING | F, 5′-tgcatccatccatcccacag-3′<br>R, 5′-caccagacaggcacttagca-3′ | Stimulator of interferon genes |
| bTBK1 | F, 5′-tgcagctactggatcactgc-3′<br>R, 5′-aacaggcatgtctccactcc-3′ | Tank binding kinase 1 |
| bIRF3 | F, 5′-aagccccacctctaaagctc-3′<br>tatcagccagggcagtatcc-3′ | Interferon regulatory factor 3 |
| bIRF7 | F, 5′-gcctcctggaaaaccaactt-3′<br>R, 5′-atcttctagggcctcgtcct-3′ | Interferon regulatory factor 7 |
| bIFN-β | F, 5′-actcctggggcagttacctt-3′<br>R, 5′-ctggtgagaatgccgaagat-3′ | Interferon-beta |
| bRPLP0 | F, 5′-cttcattgtgggagcagaca-3′<br>R, 5′-ggcaacagtttctccagagc-3′ | 60S acidic ribosomal protein large |
| bGAPDH | F, 5′-atctctgcaccttctgccga-3′<br>R, 5′-gcaggaggcattgctgaca-3′ | glyceraldehyde-3-phosphate dehydrogenase |
| *b, Bovine. | | |
